# Visuo-proprioceptive control of the hand in older adults

**DOI:** 10.1101/2020.01.18.911354

**Authors:** Hannah J. Block, Brandon M. Sexton

## Abstract

To control hand movement, we have both vision and proprioception, or position sense. The brain is known to integrate these to reduce variance. Here we ask whether older adults integrate vision and proprioception in a way that minimizes variance as young adults do, and whether older subjects compensate for an imposed visuo-proprioceptive mismatch as young adults do. Ten healthy older adults (mean age 69) and 10 healthy younger adults (mean age 19) participated. Subjects were asked to estimate the position of visual, proprioceptive, and combined targets, with no direct vision of either hand. After a veridical baseline block, a spatial visuo-proprioceptive misalignment was gradually imposed by shifting the visual component forward from the proprioceptive component without the subject’s awareness. Older subjects were more variable than young subjects at estimating both visual and proprioceptive target positions (F_1,18_ = 6.14, p = 0.023). Older subjects tended to rely more heavily on vision than proprioception compared to younger subjects. However, the weighting of vision vs. proprioception was correlated with minimum variance predictions for both older (r = 0.71, p = 0.021) and younger (r = 0.81, p = 0.0047) adults, suggesting that variance-minimizing mechanisms are present to some degree in older adults. Visual and proprioceptive realignment were similar for young and older subjects in the misalignment block, suggesting older subjects are able to realign as much as young subjects. These results suggest that intact multisensory processing in older adults should be explored as a potential means of mitigating degradation in individual sensory systems.

## Introduction

Many computations in human perception involve combining information from multiple sensory modalities. For example, in speech perception, visual information about the speaker’s mouth movements is combined with the auditory signal. Speech perception is enhanced by this process because the integrated audiovisual signal has lower variance than either unimodal signal alone (Setti et al., 2014). Similarly, for controlling movements of the hand, perception of the hand’s position in space is enhanced by integrating the visual estimate of hand position with the proprioceptive estimate (Ghahramani et al., 1997). If Ŷ_V_ and Ŷ_P_ are the brain’s visual and proprioceptive estimates of hand position, respectively, the integrated multisensory estimate, Ŷ_VP_, can be described as:

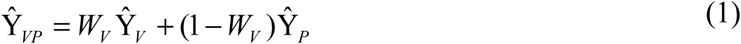

where *W_V_* is the weight of vision vs. proprioception (*W_V_* > 0.5 implies greater reliance on vision). For hand position estimation, as with speech perception, balance, and many other perceptual computations, multisensory integration is thought to follow a minimum variance model. In other words, each sensory modality is weighted in inverse proportion to its variance (Block and Bastian, 2010; Ernst and Banks, 2002; Ernst and Bulthoff, 2004; Ghahramani et al., 1997), such that the integrated percept is lower in variance than any of the unimodal percepts. Thus, multisensory integration is generally considered beneficial to human behavior. Multisensory integration gives us the flexibility to cope with the frequent sensory perturbations we experience, e.g., up-weighting proprioception when lighting is low (Mon-Williams et al., 1997); or realigning one or both sensory estimates when they become spatially misaligned (Bedford, 1999; Sober and Sabes, 2005, 2003; van Beers et al., 2002), as when washing dishes with the hands immersed in water, which refracts light.

Over the longer term, multisensory integration theoretically enables us to get the maximal benefit from sensory modalities that may worsen with age. Proprioceptive deficits have been found in the finger joint, wrist, and elbow of older adults (Adamo et al., 2012; Hughes et al., 2015), and visual deficits are well documented (Kalina, 1997). Changes in peripheral and central processing of sensory signals result in perceptual deficits that, along with impairments of certain cognitive and motor processes, cause widespread functional challenges in older adults (Mozolic et al., 2012). Hand dexterity, motor speed, and motor adaptation have all been found to decrease with age (Birren and Fisher, 1995; Liu et al., 2017; Vandevoorde and Orban de Xivry, 2019). Deficits like these can affect older adults’ capability to perform tasks of daily living, like cooking and self care, which may reduce their ability to live independently.

Given these pervasive changes to sensory systems with aging, it is important to establish the status of multisensory integration in older adults. If the computational processes underlying multisensory integration are impaired in older adults, then integration might not occur at all, or perhaps a less-reliable sensory modality is weighted too heavily, which could contribute to functional deficits. In this case, older adults could benefit from training in multisensory integration (Setti et al., 2014). Alternatively, if multisensory integration is intact, then any rehabilitation might be better spent on improving the component sensory modalities.

Unfortunately, the reality of multisensory integration in older adults appears much more complex than this simple dichotomy (de Dieuleveult et al., 2017; Mozolic et al., 2012). In some ways, older adults can appear better than young adults at multisensory integration (Mozolic et al., 2012). Older adults had equal or greater audiovisual integration in the McGurk illusion (Cienkowski and Carney, 2002) and greater response time benefits from multisensory vs. unisensory audiovisual signals (Laurienti et al., 2006). Increased integration of somatosensory signals into multisensory representation of body orientation has also been reported (Strupp et al., 1999). However, other studies have suggested that older adults are impaired at minimizing variance in multisensory integration when cognitive demands are higher; e.g., dual task, distractors, experimentally degraded sensory signals (de Dieuleveult et al., 2017). This was the case across many tasks and combinations of visual, auditory, vestibular, and somatosensory modalities (de Dieuleveult et al., 2017).

Here we investigated multisensory integration in older adults in the context of visuo-proprioceptive estimation of hand position. Proprioception plays a critical role in accurate motor function (Findlater and Dukelow, 2017) and has been found to worsen with aging (Hughes et al., 2015), but little is known about the effect of aging on visuo-proprioceptive integration. We specifically asked whether weighting of vision vs. proprioception is similar in younger and older adults, and whether integration minimizes variance equally well in these two groups. We also asked whether older adults compensate for a visuo-proprioceptive perturbation as well as younger adults do. The perturbation was imposed gradually, without subjects’ awareness, by creating a spatial mismatch between visual and proprioceptive signals about hand position.

## Methods

### Subjects

10 healthy older adults (mean 68.5 years, range 60 to 81 years) and 10 healthy younger adults (mean 19.4 years, range 18 to 21 years) were recruited. Subjects reported that they had no history of neurological disease, no upper limb musculoskeletal injuries, that they had normal or corrected to normal vision, and that they were right handed. All procedures were approved by the Indiana University institutional review board and subjects gave written informed consent.

### Task

Visuo-proprioceptive estimates of hand position are most commonly measured and/or perturbed with a bimanual task, using an “indicator” hand to indicate the subject’s perception of the “target” hand’s position when visual, proprioceptive, or both types of information about the target are available (Smeets et al., 2006; van Beers et al., 2002, 1999, 1998, 1996). Multisensory integration may differ if the goal is perception vs. action (Patterson et al., 2017), so it is crucial that sensory outcome measures are taken in the context of interest, i.e., action planning. The bimanual task is well suited to assessing multisensory perception in the context of action planning because integration of visual and proprioceptive information about the target hand occurs while subjects are planning a movement of the indicator hand.

#### Apparatus

Subjects were seated at a custom 2D reflected rear projection apparatus (Fig. 1A). The task was presented via a mirror that made it appear that the image was in the plane of the touchscreens, where the subject’s unseen hands were located. The apparatus consisted of two touch overlays (PQLabs) with a 3mm-thick pane of glass sandwiched between them. Overlays use infrared beams to detect touch on either side of the glass with < 0.5mm resolution. The indicator hand remained above the touchscreen glass throughout the experiment, and the target hand remained below the glass. A fabric drape around the shoulders prevented subjects from viewing their upper arms or shoulders, and the apparatus mirror prevented them from viewing either hand.

**Figure 1.**
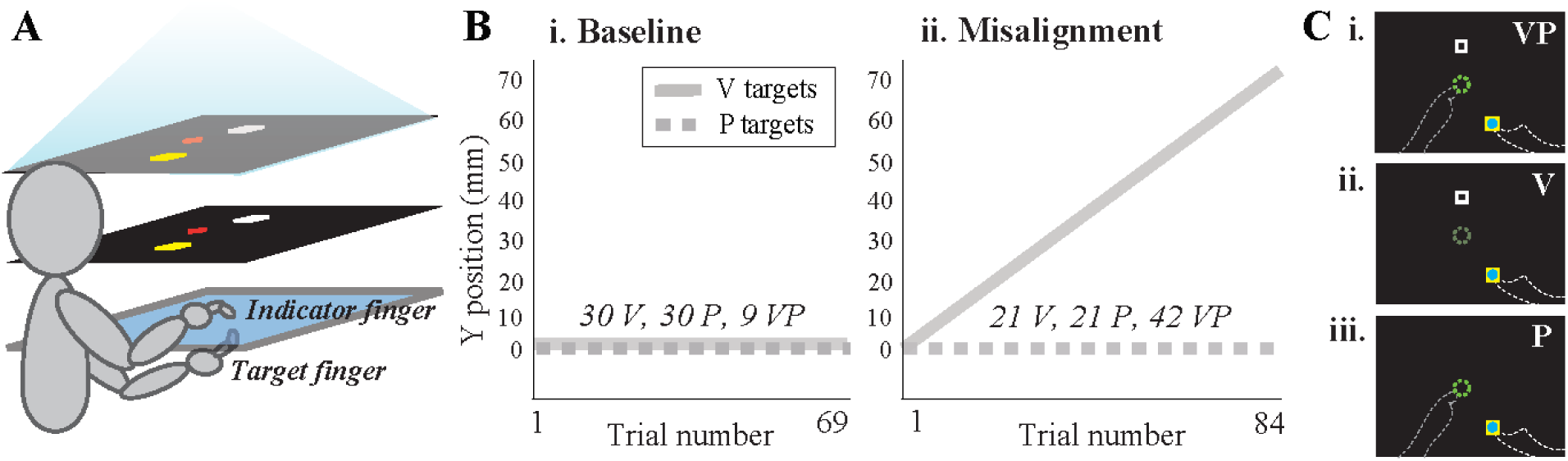
**Experiment design. A. Apparatus.** Subjects were seated at a 2D reflected rear projection apparatus. Subjects viewed the task display (top layer) in a mirror (middle layer). Subjects used their indicator finger (right hand) to indicate on top of the touchscreen glass (bottom layer) where they felt the target to be. Targets were either proprioceptive (P: left index finger positioned beneath the glass), visual (V: a white square), or both (VP: target finger with white square). The mirror was positioned equidistant from the projection screen and touchscreen glass, creating the illusion that the task display was positioned in the plane of the touchscreen. No direct vision of the hands or arms was possible. **B. Trial sequence.** Subjects performed two blocks of trials, each containing a combination of V, P and VP trials in alternating order. **i.** There was no perturbation in the baseline block, with V targets (grey line) projected veridically above P targets (dashed grey line). **ii.** In the misalignment block, the V target was gradually shifted forward from the P target to a max of 70 mm. The y-dimension was in the subject’s sagittal plane. **C. Misaligned targets.** Bird’s eye view of task display late in the misalignment block for VP (i), V (ii), and P (iii) targets. Dashed lines not visible to subject. Yellow square reflects starting position for the indicator finger. Green dashed circle reflects tactile marker where target finger was positioned for P and VP targets. During V trials, subjects rested their target hand in their lap. Subjects received no performance feedback or knowledge of results.

#### Single trial design

Subjects were instructed to move their unseen indicator finger above the glass from a start position indicated by a 1 cm yellow square to indicate where they perceived the target to be (Fig. 1A). There were three target types: VP targets (target index fingertip positioned on tactile marker beneath glass, with 1 cm white square to indicate the location visually), P targets (target finger only), and V targets (white square only).

There were five possible locations of the yellow start square, centered with the subject’s midline. An 8 mm blue cursor was shown at the indicator finger’s position only near the yellow start box, to help the subject achieve the start position and reduce any drift that might affect the indicator finger (Patterson et al., 2017). There were two possible target locations about 15 cm in front of the yellow start positions and slightly to the left of the subject’s midline. Subjects were asked to fixate a red cross that appeared randomly within a 10 cm zone centered at the subject’s midline. Start and target positions were randomly sequenced to prevent subjects from memorizing a particular indicator hand movement direction or extent. Subjects were asked to lift the indicator finger off the glass and place it down where they thought the target was, without dragging their finger along the glass. Adjustments were permitted, and the trial ended when the indicator finger did not move more than 2 mm in 2 seconds. The indicator finger’s position at this time was recorded as the endpoint. The blue cursor disappeared as soon as the indicator finger left the yellow start box, and did not reappear until the beginning of the next trial. Thus, subjects received no online or endpoint visual feedback about the indicator finger’s movements to the estimated target position.

#### Task blocks

The experiment consisted of two blocks (Fig. 1B). The <u>baseline</u> block (30 V, 30 P, and 9 VP targets in alternating order) was used to assess subjects’ variance at estimating V and P target positions, and their weighting of vision vs. proprioception when both were available (VP trials) and in the absence of a perturbation. The white box was projected veridically above the target finger on VP trials. During the <u>misalignment</u> block (42 VP, 21 V, 21 P targets in alternating order), visuo-proprioceptive misalignment in the sagittal plane was imposed gradually by shifting the white square 1.67 mm forward on each VP trial. Participants do not generally notice this change, which results in a 70 mm visuo-proprio misalignment by the end of the block (Block et al., 2013; Block and Bastian, 2012, 2011) (Fig. 1C). The entire behavioral task, including 8 practice trials that were not analyzed, took about 45 minutes.

### Data analysis

#### Variance and bias

After subtracting true target position, we computed the 2D variance of the indicator finger endpoints on the 30 V trials and 30 P trials of the baseline block. We computed spatial bias for V and P targets in baseline as the distance between true target position and the mean of the 30 V estimates and 30 P estimates, respectively.

#### Weight of vision vs. proprioception (w_V_)

We computed an experimental estimate of subjects’ weight of vision relative to proprioception (*w_V_*) on baseline VP trials:

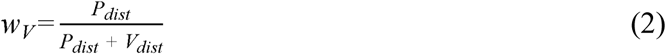

where *P_dist_* and *V_dist_* are the sagittal-plane (y-dimension) distances between the mean final position of the indicator finger on P or V targets, respectively, and the mean position of the indicator finger on VP targets (Block et al., 2013; Block and Bastian, 2012, 2011, 2010). This method takes advantage of the naturally different spatial biases inherent in the alignment of vision and proprioception (Crowe et al., 1987; Foley and Held, 1972), even with no perturbation (Smeets et al., 2006). In other words, if on VP trials a subject points closer to his P target estimate than his V target estimate, we reason that he was relying more on proprioception. Weight of vision vs. proprioception is known to differ across spatial dimensions, for example favoring proprioception in depth and vision in azimuth (van Beers et al., 2002, 1999, 1996). Because the perturbation in the misalignment block is imposed in the sagittal dimension, we chose to compute *w_V_* in that same dimension.

We also computed a predicted value of *w_V_* for each subject, based on the minimum variance model. This model, which has been supported in this and other tasks (Block and Bastian, 2010; Ernst and Banks, 2002; Ghahramani et al., 1997), predicts each sensory modality will be weighted in inverse proportion to its variance (Block and Bastian, 2010). We do not have access to the variance of the sensory modalities themselves, as the variance of subjects’ V and P target estimates reflects not only the visual or proprioceptive variance associated with the target, but also the proprioceptive variance associated with the indicator hand. However, we have previously approximated the sensory variance by assuming proprioceptive variance of the two hands is equal, and subtracting half the P target variance from both P and V target variance (Block and Bastian, 2010), and we used this method here as well:

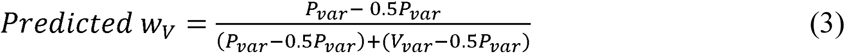

where *P_var_* and *V_var_* are the variances of the subject’s baseline P target and V target estimates in the sagittal dimension, respectively.

#### Visuo-proprioceptive realignment

In the misalignment block, VP trials were used to create the misalignment while V and P trials were used to assess visual and proprioceptive realignment. Thus, realignment measures are based on V and P trials (Fig. 1B) (Block et al., 2013; Block and Bastian, 2012, 2011). If the proprioceptive estimate of the target finger’s position, as shown by indicator finger endpoints, moves forward to close the visuo-proprioceptive gap (*Δŷ_P_*), then we expect to observe overshoot on P targets. Similarly, if perceived position of the white white square moves closer to the target finger (*Δŷ_V_*), then we expect to observe undershoot on V targets. We quantified visual and proprioceptive realignment (*Δŷ_V_* and *Δŷ_P_*) as previously (Block et al., 2013; Block and Bastian, 2012, 2011): after calculating mean indicator finger endpoint positions in the sagittal dimension on the first and last 4 V and P trials, respectively, we computed the difference relative to actual target position, which is constant for P targets but shifts 70mm for V targets:

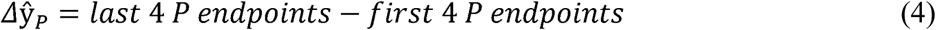

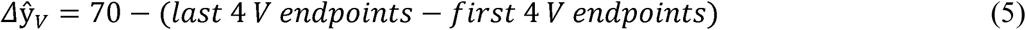

#### Statistical analysis

For each of variance, bias, and realignment, we performed a mixed model ANOVA with between-subjects factor Group (older and younger) and with-subjects factor Modality (visual and proprioceptive). We performed a two-sample t-test to compare experimental *w_V_* across groups, and computed the correlation coefficient for each group’s predicted vs. experimental *w_V_*. All hypothesis tests were performed two-tailed, with α of 0.05.

## Results

Both older and young subjects were able to perform the target estimation task. Some subjects, such as the young subject in Fig. 2A, were able to estimate V and P target positions with high precision relative to others, such as the older subject in Fig. 2C. As we have seen previously (Liu et al., 2018), all subjects exhibited some degree of spatial bias in their estimates, even the more precise subjects (Fig. 2A).

**Figure 2.**
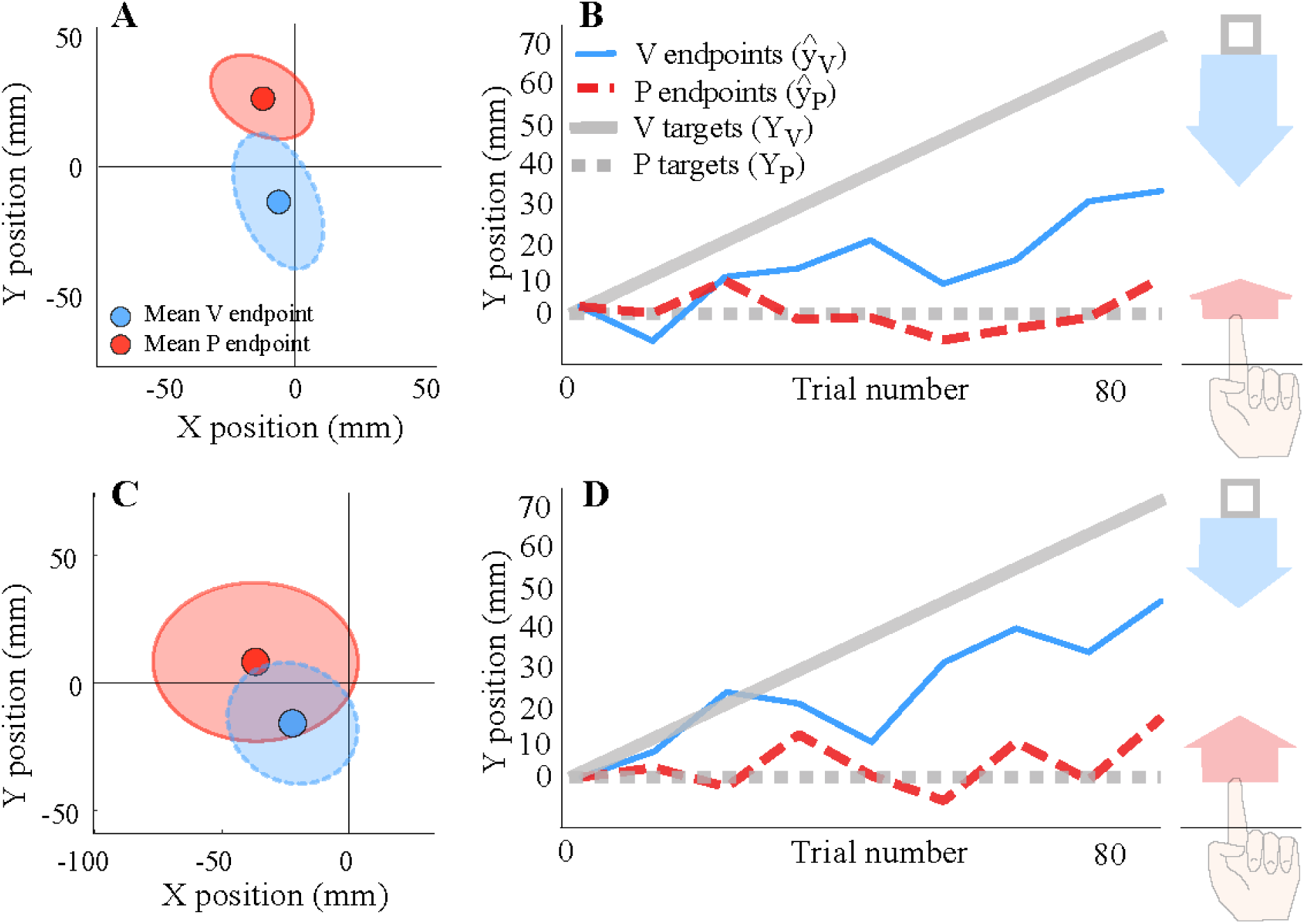
**Example young (A-B) and older (C-D) subjects. A.** Bird’s eye view of a young subject’s (age 19) baseline estimates of V and P target locations, which were always at the origin. Proprioceptive target estimates (red) were biased slightly ahead and to the left of true target position, with a 2D variance of 35.5 mm^2^. Visual target estimates (blue) were biased slightly toward the subject, with a 2D variance of 23.6 mm^2^. Shaded regions represent 90% confidence intervals. **B.** Performance of the same young subject during misalignment block. As the V target (solid grey line) shifted forward from the P target (dashed grey line), P endpoints (dashed red line) began to overshoot P targets, indicating a forward shift in perceived target position (proprioceptive realignment, red arrow) of 7.3 mm by the end of the block. At the same time, V endpoints (blue line) began to undershoot V targets, indicating a shift toward the subject in perceived position of the white square (visual realignment, blue arrow) of 40.4 mm by the end of the block. In other words, the subject compensated for 68% of the 70 mm perturbation. **C.** Bird’s eye view of an older subject’s (age 69) baseline estimates of V and P target locations, which were always at the origin. Estimates of both target locations were biased to the left, with 2D variance of 60.5 mm^2^ for P target estimates and 41.6 mm^2^ for V target estimates. **D.** Performance of the same older subject during misalignment block. The subject realigned proprioception by 15.8 mm and vision by 23.2 mm. In other words, the subject compensated for 56% of the 70 mm perturbation.

In the group analysis of target estimation variance for V and P targets during the baseline block (Fig. 3A), there was no main effect of target modality (F_1,18_ = 1.29, p = 0.27). In other words, for both young and older subjects, variance in target estimation was similar for visual vs. proprioceptive targets. There was a main effect of group (F_1,18_ = 6.14, p = 0.023), but no group x modality interaction (F_1,18_ = 1.02, p = 0.33). This suggests that older subjects estimated target positions with greater variance than young subjects, regardless of the target modality. In the group analysis of spatial bias in estimating V and P targets during the baseline block (Fig. 3B), there was no effect of group (F_1,18_ = 0.21, p = 0.65) or target modality (F_1,18_ = 0.09, p = 0.76) and no interaction (F_1,18_ = 0.04, p = 0.84).

**Figure 3.**
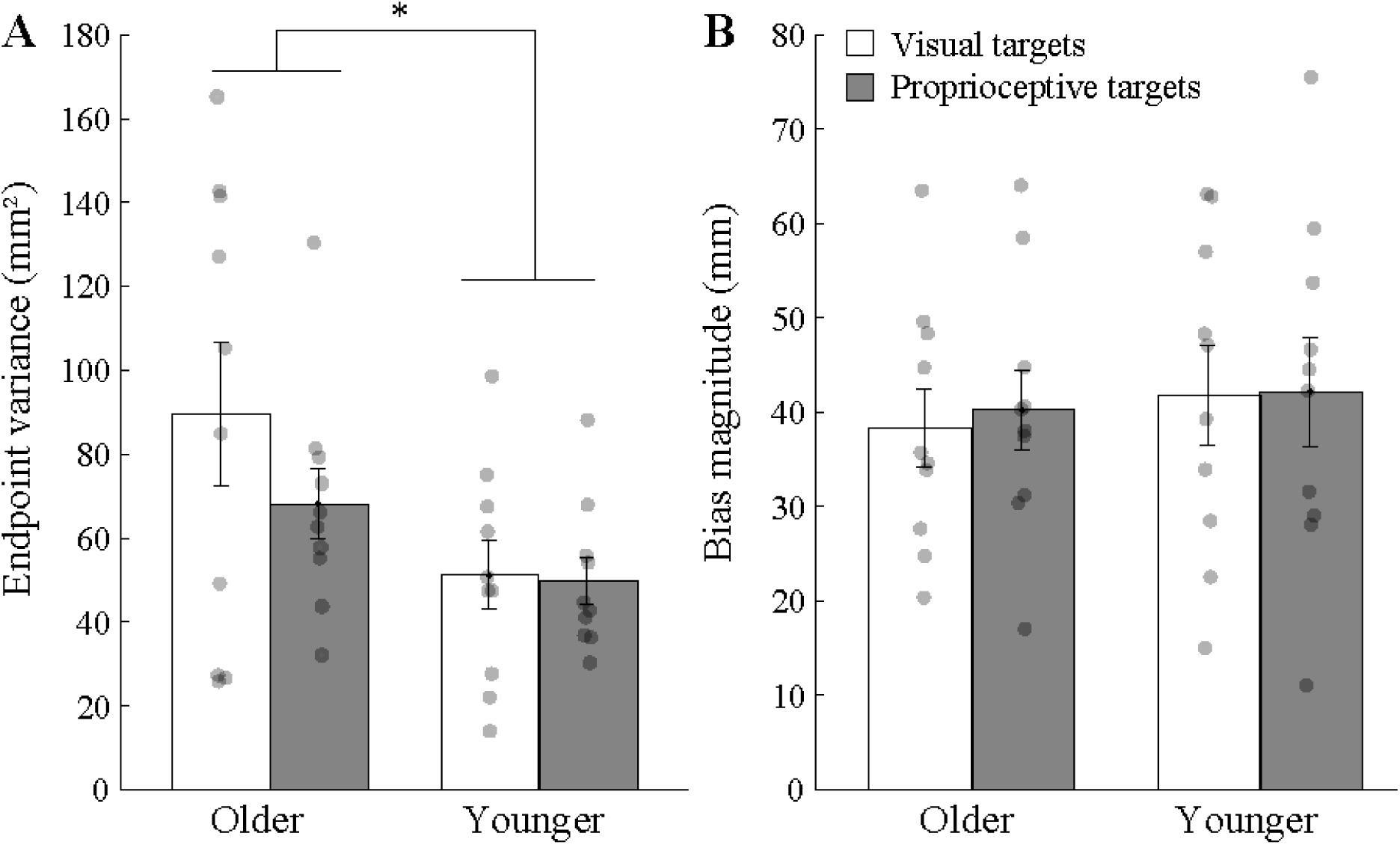
Group performance on V and P targets. **A.** Mean variance in target position estimation, by group and target modality. Older subjects estimated target positions with greater variability than younger subjects did, regardless of whether they were estimating visual (white) or proprioceptive (grey) targets. *Main effect of group, p < 0.05. **B.** Mean absolute bias in target position estimation, by group and target estimation. Both older and younger subjects were spatially biased in their estimates of V and P target locations, but bias magnitude did not differ across groups or target modalities. Error bars represent SEM. Grey circles represent individual subjects.

Weighting of vision vs. proprioception (*w_V_*) is known to differ in the sagittal vs. lateral dimensions (van Beers et al., 2002). We therefore evaluated this parameter in the sagittal or y-dimension during baseline, as this was the dimension of perturbation during the misalignment block. In our subjects, *w_V_* ranged from 0.12 (relying heavily on proprioception) to 0.86 (relying heavily on vision). Mean *w_V_* was higher in the older group than the young group (Fig. 4A), which could suggest the older subjects relied more heavily on vision, but this difference did not quite reach significance (t_18_ = 1.97, p = 0.065).

**Figure 4.**
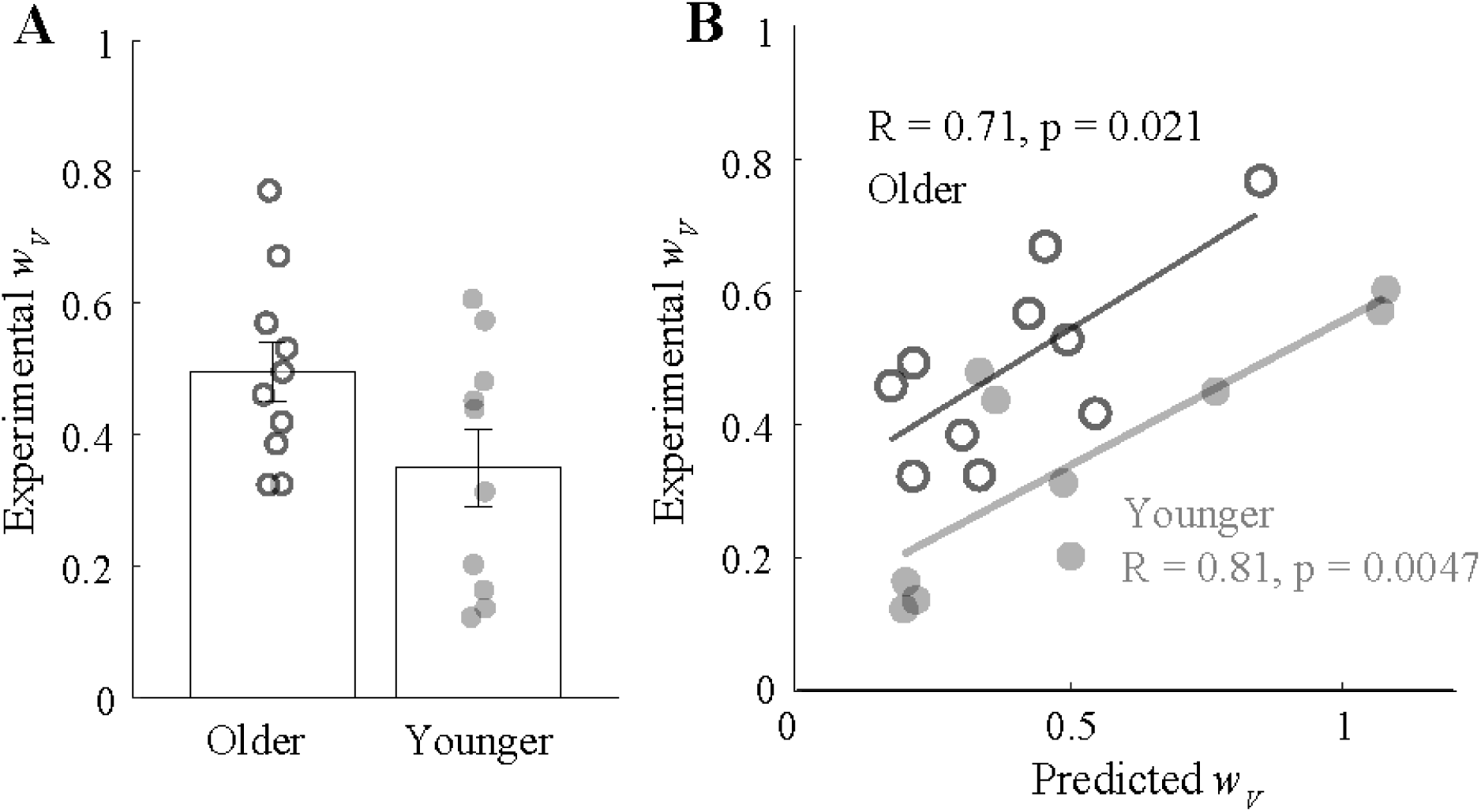
Weight of vision vs. proprioception (*w_V_*). Smaller values indicate greater reliance on proprioception while larger values indicate greater reliance on vision. **A.** Mean experimental *w_V_* did not differ significantly between young and older subjects. Error bars represent SEM. **B.** Experimental *w_V_* was correlated with the *w_V_* predicted by the minimum variance model for both older (open circles) and younger (grey circles) subjects.

This experimentally-computed value of *w_V_* relies on whether a subject’s VP target estimates are spatially closer to their V target estimates (higher *w_V_*) or to their P target estimates (lower *w_V_*); we compared experimental *w_V_* with the value that would be predicted by the minimum variance model (Ernst and Banks, 2002; Ghahramani et al., 1997). According to this model, *w_V_* should minimize variance by relying more heavily on the less-variable sensory modality. In both older and young subjects, experimental *w_V_* and predicted *w_V_* were correlated (older R = 0.71, p = 0.021; young R = 0.81, p = 0.0047); this suggests that both groups integrated vision and proprioception in a way that approximates the minimum variance model (Fig. 4B).

During the misalignment block, V targets were gradually displaced forward from P targets, creating a spatial mismatch in VP targets. None of the subjects reported a conscious awareness of the white box being displaced forward from the target finger. All subjects used some combination of visual and proprioceptive realignment to compensate for some portion of the 70 mm spatial mismatch in VP targets. The total amount of mismatch subjects compensated for ranged from 24 mm (34% of 70 mm) to 73 mm (104%). Both subjects illustrated in Fig. 2 fell in between these extremes; the young subject realigned about 68% in total, mostly by realigning their visual estimates (Fig. 2B). The older example subject realigned 56% in total, with realignment magnitude more evenly divided between vision and proprioception (Fig. 2D). In other words, subjects exhibited a wide range of responses to the misalignment perturbation, as should be expected in the absence of any performance feedback. At the group level, a main effect of modality (F_1,18_ = 6.78, p = 0.018) indicates that subjects in both groups realigned their visual estimates more than their proprioceptive estimates (Fig. 5). Based on the absence of a group effect (F_1,18_ = 1.83, p = 0.19) or interaction (F_1,18_ = 0.87, p = 0.36), this data does not support a relationship between age and visuo-proprioceptive realignment.

**Figure 5.**
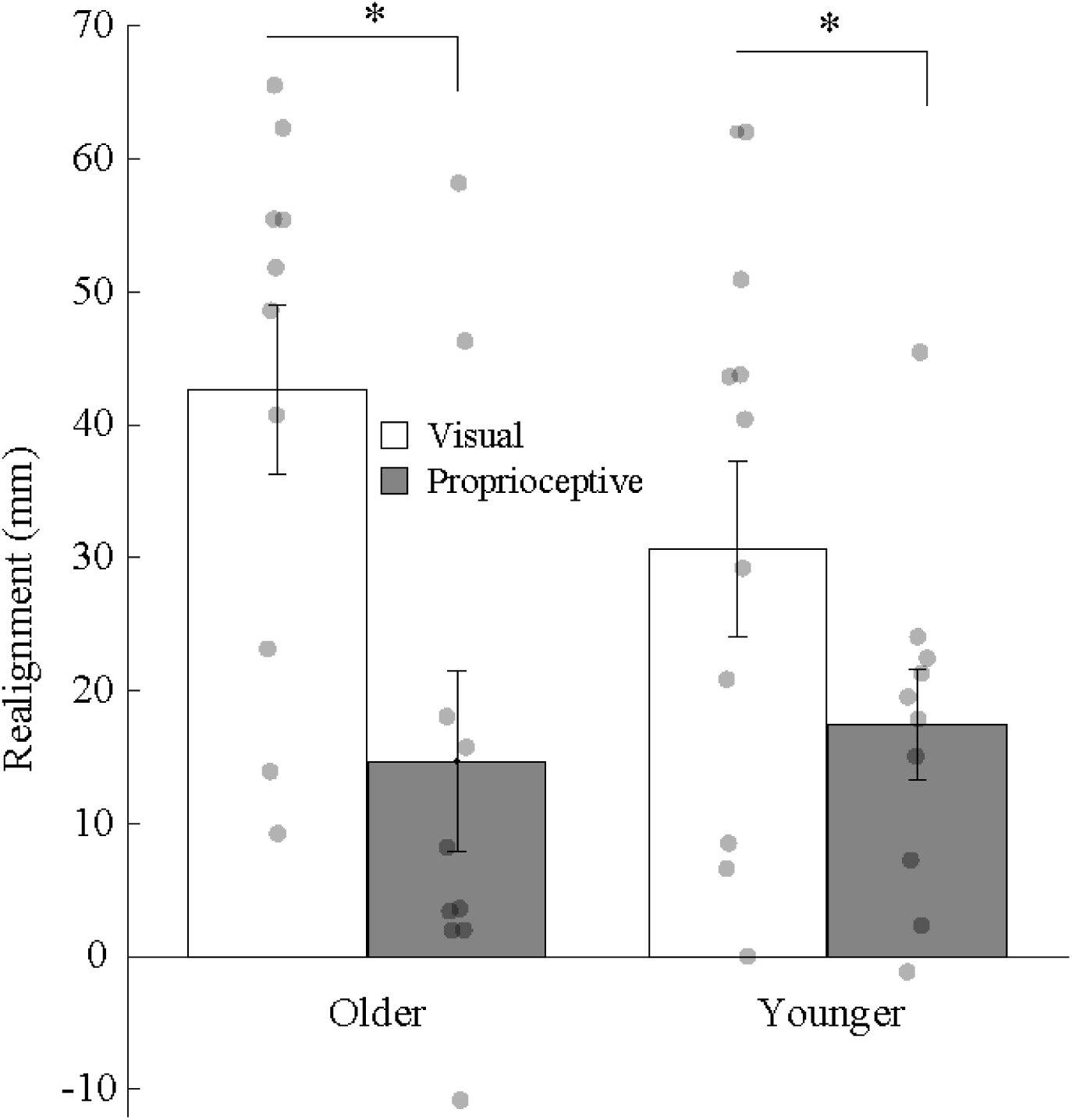
Mean realignment by group and modality. Younger and older subjects realigned vision and proprioception similarly. Both groups realigned vision a greater magnitude than proprioception. *Main effect of realignment modality, p < 0.05. Error bars represent SEM. Grey circles represent individual subjects.

At the conclusion of each session, subjects were asked to rate on a scale of 1-10 their attention level and fatigue. Ratings provided by subjects revealed there was no difference in perceived attention (t_18_ = 1.23, p = 0.24) or fatigue (t_18_ = 0.17, p = 0.87) between older and younger subjects.

## Discussion

These results suggest that older subjects are more variable when pointing to visual and proprioceptive targets than younger subjects. Older adults also tended to rely more heavily on vision when both modalities were present, although their visuo-proprioceptive weightings were correlated with minimum variance predictions. Younger and older adults realigned vision and proprioception similarly in response to a visuo-proprioceptive mismatch. Taken together, these findings support intact multisensory processing of vision and proprioception in older adults.

### Variance and bias

In our subjects, older adults were more variable but not more biased than younger adults at indicating the position of visual and proprioceptive targets, and there was no effect of target modality. Increased variance with age is consistent with the literature on declining sensory perception (Adamo et al., 2012; Hughes et al., 2015; Kalina, 1997). Older adults had greater variance in visual and vestibular perception of vertical orientation, and perception was more easily affected by changes in the visual cue (Alberts et al., 2019). Perceptual acuity is usually defined in terms of variance rather than systematic bias, making it difficult to know whether to expect larger biases in older adults. We have previously found worsening of proprioceptive bias with age in a tablet-based task (Hoseini et al., 2015); however, that study was substantially larger. Thus, age-related bias differences cannot be ruled out by the absence of a difference with our present sample size.

One important question to consider in a perceptual estimation task involving movement is whether the greater endpoint variance of older adults could be due to more variable motor control rather than sensory perception. While this is a possibility, it must be remembered that movements of the indicator finger had no time constraints or limits on adjustments. Instead, subjects were told to move at a comfortable pace, to be as accurate as possible, and to make adjustments if needed. Another possibility is that older adults were more variable because of greater fatigue. However, the fatigue ratings did not differ across groups, and variance was assessed from the baseline block, which was early in the session, so we do not think this is a likely explanation. Indeed, the task was originally designed to be comfortable for patients with moderate to severe sensorimotor deficits (Block and Bastian, 2012); subjects rested their right elbow on the apparatus and their left hand in their lap other than for proprioceptive targets, when they were asked to touch the tactile marker for several seconds.

### Weight of vision vs. proprioception

For both younger and older adults, experimentally-computed weighting of vision vs. proprioception was correlated with the minimum variance model prediction. Correlations in small sample sizes must be interpreted with caution, but these results are consistent with some mechanism of minimizing variance in visuo-proprioceptive integration at work in both older and younger adults. In our subjects, there was also a tendency for older adults to rely on vision more heavily than proprioception, compared to younger adults. This is consistent with studies of visuo-vestibular perception of the vertical in older adults (Alberts et al., 2019; Kobayashi et al., 2002; Sun et al., 2014). In brightly lit surroundings, we might expect most people to favor vision over proprioception. However, because the task is performed in a dark room with a sparse visual display, reliance on proprioception can equal or exceed reliance on vision (Mon-Williams et al., 1997). Further study with a larger sample is likely needed to establish whether older adults are indeed more likely to favor vision than younger adults, as suggested by our data. This would not be a surprising result: both vision and proprioception are known to degrade with age; however, unlike proprioception, vision can be improved by corrective lenses. This being the case, we might wonder why older subjects had greater variance at estimating visual as well as proprioceptive targets. It is possible that impaired proprioception in the indicator hand contributed to greater overall variance in older adults’ target estimations; a non-bimanual method of assessing subjects’ perception could clarify this question.

An alternative interpretation of our results is that older adults did not integrate vision and proprioception in a way that the minimum variance model would predict. In other words, if the older adults really did have similarly variable vision and proprioception, their greater reliance on vision would be non-optimal. At the same time, the correlation between experimental and minimum variance-predicted weighting in older adults suggest there is some influence of a variance-minimizing process. In other words, our weighting results are best explained as a combination of minimum variance integration with other processes that are affected by aging. Hirst et al. (2019) reached a similar conclusion with the sound-flash illusion, a measure of auditory-visual integration; they found the effect to be influenced, but not fully explained, by age-related increases in auditory and visual variance (Hirst et al., 2019).

### Visuo-proprioceptive realignment

Older and younger subjects realigned similarly in response to a gradually-imposed 70 mm mismatch of vision and proprioception. This is consistent with the absence of an age-related impairment in multisensory integration. Clearly, we could not conclude that visuo-proprioceptive realignment is unaffected by aging from such a small study. However, the fact that variance differences could be detected in this sample suggests we had the power to detect multisensory differences between groups, had they been as robust as the variance differences. We therefore suggest only that these aspects of multisensory processing may be less impaired by aging than unisensory variance is. There is very limited literature on spatial recalibration of sensory modalities in older adults, but it appears consistent with our finding of similar realignment in older and younger adults. Motor adaptation of reaching is a cerebellum-dependent process which likely involves proprioceptive realignment (Mostafa et al., 2015; Ostry and Gribble, 2016). While motor adaptation declines with age, this appears largely due to deficits in the cognitive aspects of this type of learning; the implicit component, internal model recalibration, is intact in older adults and may compensate for cognitive deficits (Vandevoorde and Orban de Xivry, 2019). More directly relevant to the present study, Cressman et al. (2010) specifically examined the proprioceptive recalibration associated with visuomotor adaptation and found that older and younger adults realigned proprioception similarly (Cressman et al., 2010). Vachon et al. (2019) found that older adults’ proprioceptive recalibration was more influenced by visual training than young adults (Vachon et al., 2019). This is consistent with the finding that older adults depend more heavily on vision, regardless of whether this would minimize variance (Alberts et al., 2019; de Dieuleveult et al., 2017; Mozolic et al., 2012).

Work is needed in many domains of perception to understand age-related changes in multisensory integration, which is not well elucidated even in young adults. Curiously, multisensory integration in older adults can be more robust in older adults than younger adults (Mozolic et al., 2012). This has been demonstrated across numerous domains, including multisensory perception of speech (Cienkowski and Carney, 2002) and body orientation (Strupp et al., 1999), and response time advantages to multisensory targets (Diederich et al., 2008; Laurienti et al., 2006; Peiffer et al., 2007). Such effects have been attributed to increased noise at baseline in older adults (de Dieuleveult et al., 2017; Mozolic et al., 2012).

However, while multisensory integration can be considered beneficial in general, these age-related changes can be both enhancements and impairments (Basharat et al., 2019). For example, in the temporal domain, older adults’ response times are more facilitated by multisensory stimuli than younger adults, which is considered a perceptual enhancement (Basharat et al., 2019). Older adults also have larger temporal binding windows, the range of time during which multisensory events can be perceived as happening at the same time; while this increases the chance of multisensory integration occurring, it also increases the chance of two signals that arise from different events being erroneously encoded as a single event (Basharat et al., 2019). This is considered a perceptual impairment, as it can reduce speech comprehension (Maguinness et al., 2011), increase fall risk (Mahoney et al., 2014), and make it difficult to ignore distracting or irrelevant information (Wu et al., 2012). In a systematic review, De Deiuleveult et al. (2017) concluded that older adults use all available sensory modalities, even distractors and disrupted information that could impair performance of the task.

### Conclusions

Here we found that older adults, while more variable at estimating visual and proprioceptive target positions than younger adults, were similarly able to compensate for a visuo-proprioceptive mismatch. Older adults may have relied on vision more than younger adults, and possibly more than warranted by visual and proprioceptive variance, but their weightings did appear to be at least influenced by variance-minimizing mechanisms. Given the many functional challenges faced by older adults as a result of peripheral and central decline in unisensory systems, the interest in multisensory integration, which appears intact in some ways, is not surprising. Much further study across perceptual domains will be needed to draw any principles that could be applied by future rehabilitation professionals.

